# Biosynthesis of dihydroquercetin in *Escherichia coli* from glycerol

**DOI:** 10.1101/2020.11.27.401000

**Authors:** Seon Young Park, Dongsoo Yang, Shin Hee Ha, Sang Yup Lee

## Abstract

Phenylpropanoids are a group of diverse natural products derived from aromatic amino acids. Although their demands are high both as drugs and nutraceuticals, their production mainly depends on inefficient extraction from plants. To achieve sustainable production of phenylpropanoids, engineering model microorganisms such *as Escherichia coli* has been sought, but most strains require supplementation of expensive precursors. Here, we report one-step production of a representative phenylpropanoid, dihydroquercetin (DHQ), from simple carbon sources in *E. coli* for the first time. The best DHQ producer strain capable of producing 239.4 mg/L of DHQ from glycerol was obtained by optimizing the biosynthetic pathway and engineering the signal peptide of cytochrome P450 (TT7) from *Arabidopsis thaliana*. The engineered plant P450 could produce a significantly higher titer of DHQ than a bacterial monooxygenase, showing the potential of employing plant P450s for the production of diverse natural products that has been previously difficult in bacterial hosts. This study will serve as a guideline for industrial production of pharmaceutically important yet complex natural products.

## 1. Introduction

Phenylpropanoids span a diverse group of plant secondary metabolites derived from aromatic amino acids (Dey, 2016). These natural products are biologically active since they perform essential functions for plant survival and reproduction such as defence against pathogens, pests, or animals and attracting insect pollinators. Therefore, many phenylpropanoids are also reported to be beneficial for human health, showing diverse pharmaceutical activities such as antioxidant, antitumor, anti-inflammatory, antidiabetic, antiviral, and anti-bacterial properties (Dey, 2016). DHQ (also known as taxifolin) possesses various pharmacological activities including anticancer, antioxidant, chemopreventive, anti-melanogenesis, and anti-Alzheimer’s effects (An et al., 2008; Lee et al., 2007; Sato et al., 2013; Sunil and Xu, 2019), and is included in a number of nutraceuticals or drugs such as Legalon^®^ or Pycnogenol^®^ (Sunil and Xu, 2019; Yang et al., 2020b). DHQ also is an important precursor for the production of more complex flavonolignans such as silybin and isosilybin which are found from the seed of Milk Thistle (*Silybum marianum*) (Bijak, 2017; Gazak et al., 2007; Mengs et al., 2012; Deep et al., 2007).

Despite rising demands for these phenylpropanoids due to their versatile applications, their supply has been limited by unstable, toxic, and inefficient extraction from the plants. Nevertheless, this inefficient process has been the sole method for commercial production of phenylpropanoids, leading to their high prices in the market. In this regard, metabolic engineering has been an attractive solution for efficient and sustainable production of valuable natural products from model microorganisms such as *Escherichia coli* and *Saccharomyces cerevisiae*. Various tools and strategies developed for metabolic engineering have enabled successful production of various important natural products by microorganisms, with the state-of-the-art examples of cannabinoids (Luo et al., 2019), tropane alkaloids (Srinivasan and Smolke, 2020), and astaxanthin (Park et al., 2018), to list a few. Microbial production of DHQ from glucose was recently achieved by *Yarrowia lypotica* (110.5 mg/L) (Lv et al., 2019a) and by *S. cerevisiae* (336.8 mg/L) (Yang et al., 2020b), but bacterial production of DHQ from simple carbon sources has not yet been achieved.

In this paper, we report one-step *de novo* production of a representative phenylpropanoid, DHQ, from glycerol by recombinant *E. coli*. An *E. coli* strain capable of producing high level of DHQ was developed and optimized by biosynthetic pathway optimization, plant P450 engineering, and fermentation condition optimization. This study will serve as an important guideline by providing essential metabolic engineering strategies for industrial production of pharmaceutically important yet complex natural products.

## 2. Materials and Methods

### 2.1. Materials and strains

Eriodictyol, dihydrokaempferol (DHK), and DHQ were purchased from Merck (Sigma Aldrich). *p*-Coumaric acid and naringenin were purchased from Tokyo Chemical Industry. *TT7* (codon optimized for *E. coli*) was synthesized as gBlocks Gene Fragment from Integrated DNA Technologies Inc. The *ATR2* gene (codon optimized for *E. coli*) was synthesized from GenScript. All strains and plasmids used in this study are listed in Table 1 and 2, respectively. *E. coli* DH5α (Invitrogen) was used for routine gene cloning works. BTY5 (Kim et al., 2018) was used as a host strain for the production of all phenylpropanoids.

**Table 1.** List of strains used in this study.

| Name | Description | Source |
| --- | --- | --- |
| DH5α | F<sup>-</sup> ϕ80 *lacZ*Δ*M15* $\Delta$(*lacZ**Y**A*-*argF*)U169 *recA1* *endA1* *hsdR17* (*r*<sub>K</sub><sup>-</sup>, *m*<sub>K</sub><sup>+</sup>) *phoA* *supE44* *thi-1* *gyrA96* *relA1* λ<sup>-</sup> | Invitrogen |
| BL21(DE3) | F<sup>-</sup> *ompT* *hsdSB* (*rB-mB*-) *gal dcm* (DE3) | Invitrogen |
| BTY5 | BL21 (DE3) Δ*tyrR* Δ*tyrP* | (Kim et al., 2018) |
| BTY5.13 | BTY5 harboring pTY13 | (Kim et al., 2018) |
| DHQ | BTY5 harboring pCOU, pDHK, and pDHQ | This study |

**Table 2.**
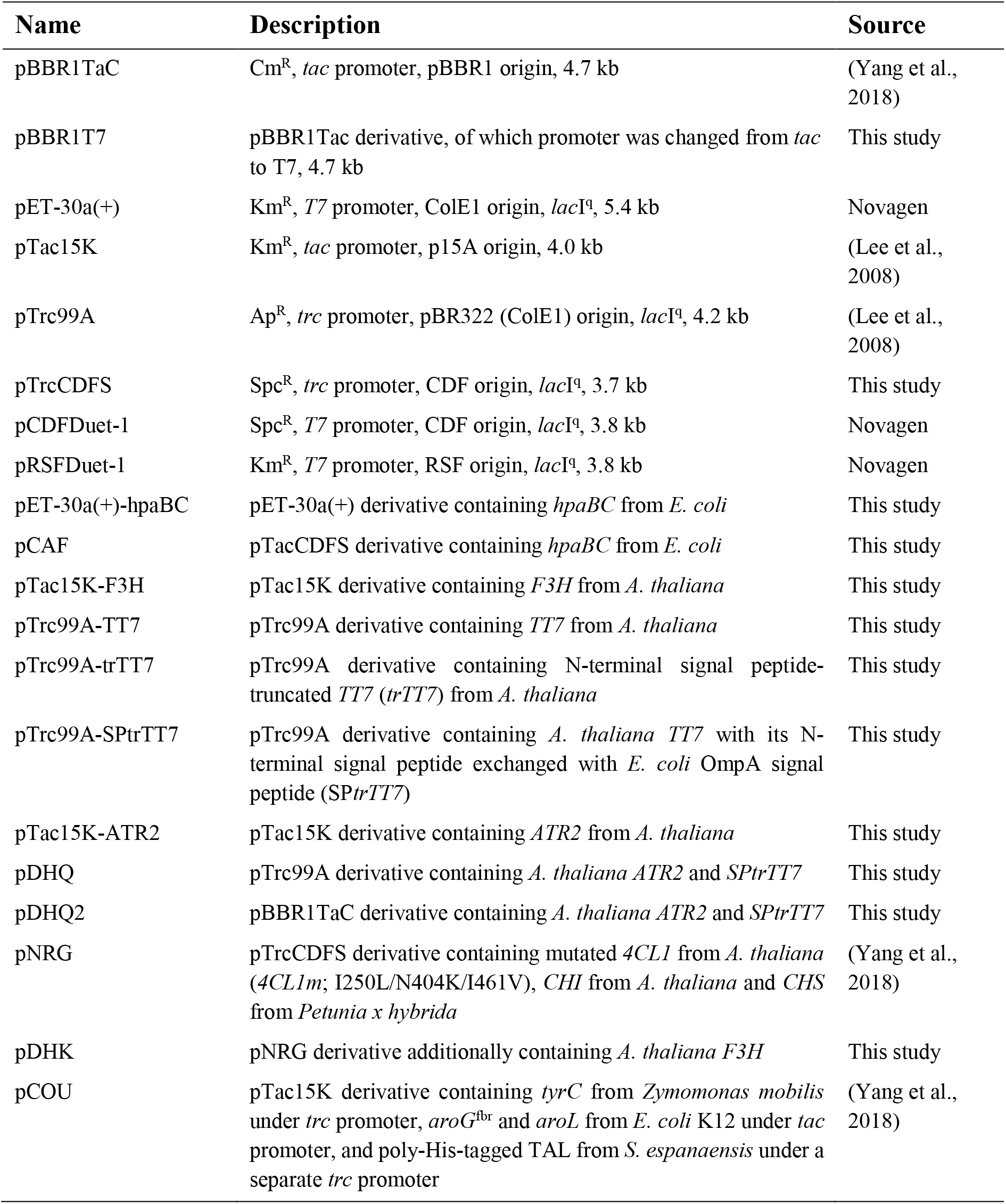
List of plasmids used in this study. Abbreviations: Ap, ampicillin; Km, kanamycin; Cm, chloramphenicol; Spc, spectinomycin; R, resistance.

### 2.2. Plasmid construction

Standard protocols were used for PCR, gel electrophoresis and transformation experiments (Sambrook, 1989). The plasmids and oligonucleotides used in this study are listed in Table 2 and 3, respectively. *E. coli* DH5α (Invitrogen) was used as a host strain for routine gene cloning, in Luria-Bertani (LB) medium (per liter: 10 g tryptone, 5 g yeast extract and 10 g NaCl) or on LB agar plates at 37 °C supplemented with appropriate concentrations of antibiotics when necessary: 50 μg/mL of kanamycin, 100 μg/mL of ampicillin, 100 μg/mL of spectinomycin, and/or 17 μg/mL of chloramphenicol. Polymerases used for PCR reactions were either Lamp-Pfu or Pfu purchased from Biofact (Daejeon, Republic of Korea). Restriction endonucleases were purchased from either Enzynomics (Daejeon, Republic of Korea) or NEB (Ipswich, MA).

**Table 3.**
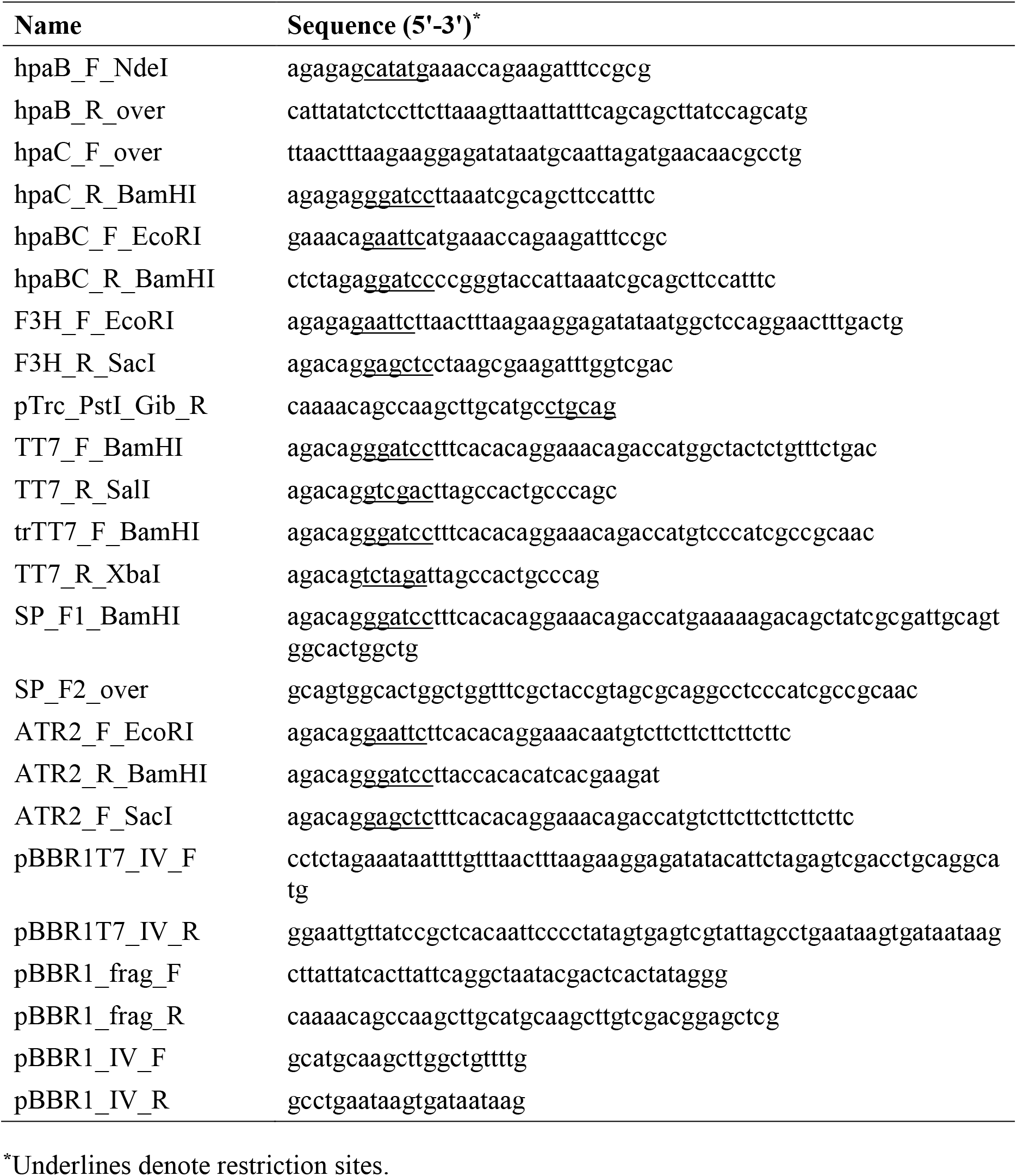
List of oligonucleotides used in this study.

To test the conversion of naringenin to DHK, plasmid pTac15K-F3H was constructed by PCR amplification of *F3H* from the cDNA of *A. thaliana* using primers F3H_F_EcoRI and F3H_R_SacI and inserting the amplified gene fragment to pTac15K at the EcoRI and SacI sites. pTrc99A-F3H was constructed using the same method. Then, the *F3H* gene was PCR amplified from pTrc99A-F3H using the primers pTrc_PstI_Gib_F and pTrc_PstI_Gib_R, and the amplified gene fragment was inserted into pNRG by Gibson assembly at PstI site to construct pDHK.

For the production of DHQ, *TT7* and *ATR2* from *A. thaliana* codon-optimized for expression in *E. coli* were synthesized (Integrated DNA Technologies Inc.). Plasmid pTrc99A-TT7 was constructed by PCR amplification of *TT7* from the artificially synthesized *TT7* gene fragment using primers TT7_F_BamHI and TT7_R_SalI followed by insertion to pTrc99A at BamHI and SalI sites. For the construction of pTrc99A-trTT7, *trTT7* was PCR amplified from the synthetic *TT7* using primers trTT7_F_BamHI and TT7_R_SalI and inserted to pTrc99A at the same sites. pTrc99A-SPtrTT7 was also constructed by PCR amplification of *SPtrTT7* from the synthetic *TT7* and inserting the amplified gene fragment to pTrc99A at the same sites. Here, *SPtrTT7* was first amplified using primers SP_F2_over and TT7_R_SalI, and then extended by PCR using primers SP_F1_BamHI and TT7_R_SalI. For the construction of pTac15K-ATR2 and pDHQ, *ATR2* was PCR amplified from the synthetic *ATR2* using primers ATR2_F_EcoRI and ATR2_R_BamHI, and was inserted to pTac15K and pTrc99A-SPtrTT7, respectively, both at EcoRI and BamHI sites. To construct pDHQ2, DNA fragment containing *ATR2* and *SPtrTT7* was PCR amplified from pDHQ using primers ATR2_F_SacI and TT7_R_XbaI, and was inserted into pBBR1TaC at SacI and XbaI sites.

### 2.3. Media and culture conditions

Strains for the production of all phenylpropanoids were first inoculated from colonies on LB agar plates into 25 mL test tubes containing 10 mL of LB medium supplemented with appropriate antibiotics, and were cultivated in a rotary shaker at 220 rpm, at 37 °C for 12-16 h. Then, 1 mL aliquot of each seed culture was transferred to a 250 mL baffled flask containing 50 mL of minimal medium (MR) and was incubated at 30 °C and at 200 rpm. For the production of DHQ, 2 g/L of yeast extract, 20 g/L of glycerol, and 3 g/L of (NH_4_)_2_SO_4_ was supplemented. The MR medium (pH 6.8) contains the followings per liter: 6.67 g KH_2_PO_4_, 4 g (NH_4_)_2_HPO_4_, 0.85 g citric acid, 0.8 g MgSO_4_·7H_2_O, and 5 mL TMS. The TMS contains the followings per liter of 5 M HCl: 10 g FeSO_4_·7H_2_O, 2.25 g ZnSO_4_·7H_2_O, 1 g CuSO_4_·5H_2_O, 0.5 g MnSO_4_·5H_2_O, 0.23 g Na_2_B_4_O_7_·10H_2_O, 2 g CaCl_2_·2H_2_O and 0.1 g (NH_4_)_6_Mo_7_O_24_ (Jeong and Lee, 2002). When the OD_600_ of the cultures reached 1–2, 1 mM isopropyl β-D-1-thiogalactopyranoside (IPTG) was added to induce heterologous gene expression. When required, 50 mg/L of kanamycin (Km), 100 mg/L of ampicillin (Ap), 17 mg/L of chloramphenicol (Cm) and/or 100 mg/L of spectinomycin (Spc) was added to the medium. After induction, the cells were cultivated for 48 hours.

Cell growth was monitored by measuring the absorbance at 600 nm (OD_600_) with an Ultrospec 3100 spectrophotometer (Amersham Biosciences, Uppsala, Sweden). The dry cell weight (DCW) was measured after drying the cell pellets at 75°C over 48 h.

### 2.4. Fed-batch fermentation

Fed-batch fermentations for DHQ production were conducted in a 5 L jar fermenter (MARADO-05D-PS, CNS, Daejeon, Republic of Korea) containing 2 L of R/2 medium supplemented with 2 g/L yeast extract, 20 g/L glycerol, and 3 g/L of (NH_4_)_2_SO_4_. When required, 50 mg/L of Km, 100 mg/L of Ap, 17 mg/L of Cm and/or 100 mg/L of Spc was added to the medium. Strains were first inoculated from colonies on LB agar plates into 25 mL test tubes each containing 10 mL of LB medium supplemented with appropriate antibiotics, and were cultivated in a rotary shaker at 220 rpm, at 37°C overnight (12-16 h). Then, 1 mL aliquot of each seed culture was transferred to a 250 mL baffled flask containing 50 mL of minimal medium supplemented with 2 g/L yeast extract, 20 g/L glycerol, and 3 g/L of (NH_4_)_2_SO_4_. After incubation at 37 °C shaken at 220 rpm until the OD_600_ value reached 3∼4, 50 mL of the seed culture was inoculated into the bioreactor containing the medium saturated by filtered air. The culture pH was controlled at 6.8 using 28% (v/v) ammonia solution (Junsei Chemical, Japan). The dissolved oxygen (DO) level of the culture was maintained at 40% of air saturation by supplying air at 2 L/min and automatically increasing the agitation speed from 200 rpm up to 1000 rpm and by changing the percentage of pure oxygen added. The pH-stat feeding strategy was employed in order to supply exhausted nutrients to the fermenter. The feeding solution contains the followings per liter: 800 g glycerol, 12 g MgSO_4_·7H_2_O, 40 g (NH_4_)_2_SO_4_, and 6 mL TMS. When pH becomes higher than 6.85 due to carbon source exhaustion, the feeding solution was automatically added. To induce heterologous gene expression, 1 mM of IPTG was added when the OD_600_ of the culture reached 20∼30.

### 2.5. Analytical procedures

After shake flask or fed-batch culture, cells were harvested by centrifugation at 16,000 *g* for 2 min. For the analysis of naringenin, eriodictyol, DHK, and DHQ, culture supernatant was extracted with an equal volume of ethyl acetate, and was vortexed vigorously using Thermo shaker (TS100, Ruicheng) for 10 min at 40 °C, 1500 rpm. The extracted ethyl acetate portion was dried and re-solubilized in methanol. The prepared samples were analyzed with high-performance liquid chromatography (HPLC; 1260 Infinity II; Agilent) equipped with DAD detectors (G7115A; Agilent) and a C18 column (Poroshell 120 EC-C18 column; 4.6 × 150 mm; Agilent). The mobile phase consists of A (0.1% trifluoroacetic acid) and solvent B (acetonitrile) was run at a flow rate of 0.6 mL/min. The following gradient was applied: 0-3 min, an isocratic condition at 5% B; 3-20 min, a linear gradient of solvent B from 5% to 70%; 20-25 min, an isocratic condition at 70% B (all in vol%). Samples were monitored at 220 nm (DHK and DHQ) or 330 nm (*p*-coumaric acid, eriodictyol, and naringenin). Concentrations of each product was determined by mapping the area of HPLC peaks to each calibration curve generated using dilutions of authentic chemicals.

DHQ produced by engineered *E. coli* strains from glycerol were further analyzed through HPLC (1100 Series HPLC; Agilent) connected with MS (LC/MSD VL; Agilent) and by comparing with the mass spectrum of the commercially available authentic standard chemicals. For DHQ, 1 mL of the culture supernatant was extracted with equal volume of ethyl acetate, dried, and re-solubilized in equal volume of methanol. XBridge C18 column (4.6 × 150 mm; Waters) was used and operated at 25 °C. For DHQ, two mobile phase solvents were used: solvent A (20 mM ammonium acetate, pH 9.6) and solvent B (acetonitrile). The total flow rate was maintained at 0.4 mL/min, and the following gradient was applied: 0-3 min, 5% solvent B; 3-15 min, a linear gradient of solvent B from 5% to 70%; 15-20 min, a linear gradient of solvent B from 70% to 90%; 20-30 min, an isocratic condition at 90% solvent B (all in vol%). The eluent was continuously injected into the mass spectrometry using ESI negative ion mode with the following conditions: fragmentor, 160 V; drying gas flow, 12.0 L/min; drying gas temperature, 350 °C; nebulizer pressure, 30 psig; capillary voltage, 5.5 kV. For analysis, scan mode was used. For LC-MS, the scanned mass range was *m/z* of 130-300.

### 2.6. SDS-PAGE analysis

To confirm the expression of heterologous enzymes, *E. coli* BL21(DE3) strains harboring the respective recombinant plasmids were first cultured in 10 mL test tubes containing R/2 medium supplemented with 2 g/L yeast extract, 20 g/L glycerol, and appropriate antibiotics. The cells were grown at 30 °C and were induced with 1 mM IPTG when the OD_600_ of the culture reached 0.4-0.6. After additional cultivation for 6-10 hr, cells were collected (i.e., 3 mL of cells when their OD_600_ reached 1.0) by centrifugation at 15,000 *g* for 1 min at 4 °C. Cell pellets were washed with 1 mL of cold phosphate-buffered saline (PBS) solution (pH 7.4), centrifuged at 15,000 *g* for 1 min at 4 °C, and resuspended in 0.3 mL of the same buffer. The resuspended cells were lysed by sonication (High-Intensity Ultrasonic Liquid Processors, Sonics & Materials Inc., Newtown, CT). To precipitate insoluble protein fractions and partially disrupted cells, the sonicated samples were centrifuged at 15,000 *g* for 10 min at 4 °C. Crude cell lysates were used for total protein expression analysis and the supernatants were used for soluble protein expression analysis.

### 2.7. Statistical analysis

We did not predetermine sample sizes. All colonies were randomly selected from plates containing ∼100–200 colonies and subject to independent flask culture and chemical analysis. All numerical data are presented as mean ± SD (standard deviation) from experiments done in duplicates or triplicates. The investigators were blinded to the group allocation by randomly selecting single colonies multiple times.

## 3. Results and discussion

DHQ is produced from *p*-coumaric acid. In *S. cerevisiae*, one-step *de novo* production of 336.8 mg/L of DHQ from glucose was reported (Lv et al., 2019a). However, in *E. coli*, production of DHQ has been achieved only by supplying expensive precursors (e.g., L-tyrosine, *p*-coumaric acid). We previously reported the construction of a recombinant *E. coli* strain capable of producing 103.8 mg/L of naringenin from glycerol (Yang et al., 2018). The naringenin producer employed tyrosine-ammonia lyase (TAL) from *S. espanaensis*, a mutant 4-coumarate:CoA ligase 1 (4CL1m) from *A. thaliana*, chalcone isomerase (CHI) from *A. thaliana*, and chalcone synthase (CHS) from *Petunia x hybrida* to enable the conversion of L-tyrosine to naringenin (Fig. 1). As naringenin is a precursor of DHQ, the naringenin producer [BTY5 harboring pCOU and pNRG (harboring *4CL1m, CHI*, and *CHS*)] was used for *de novo* DHQ production.

**Fig. 1.**
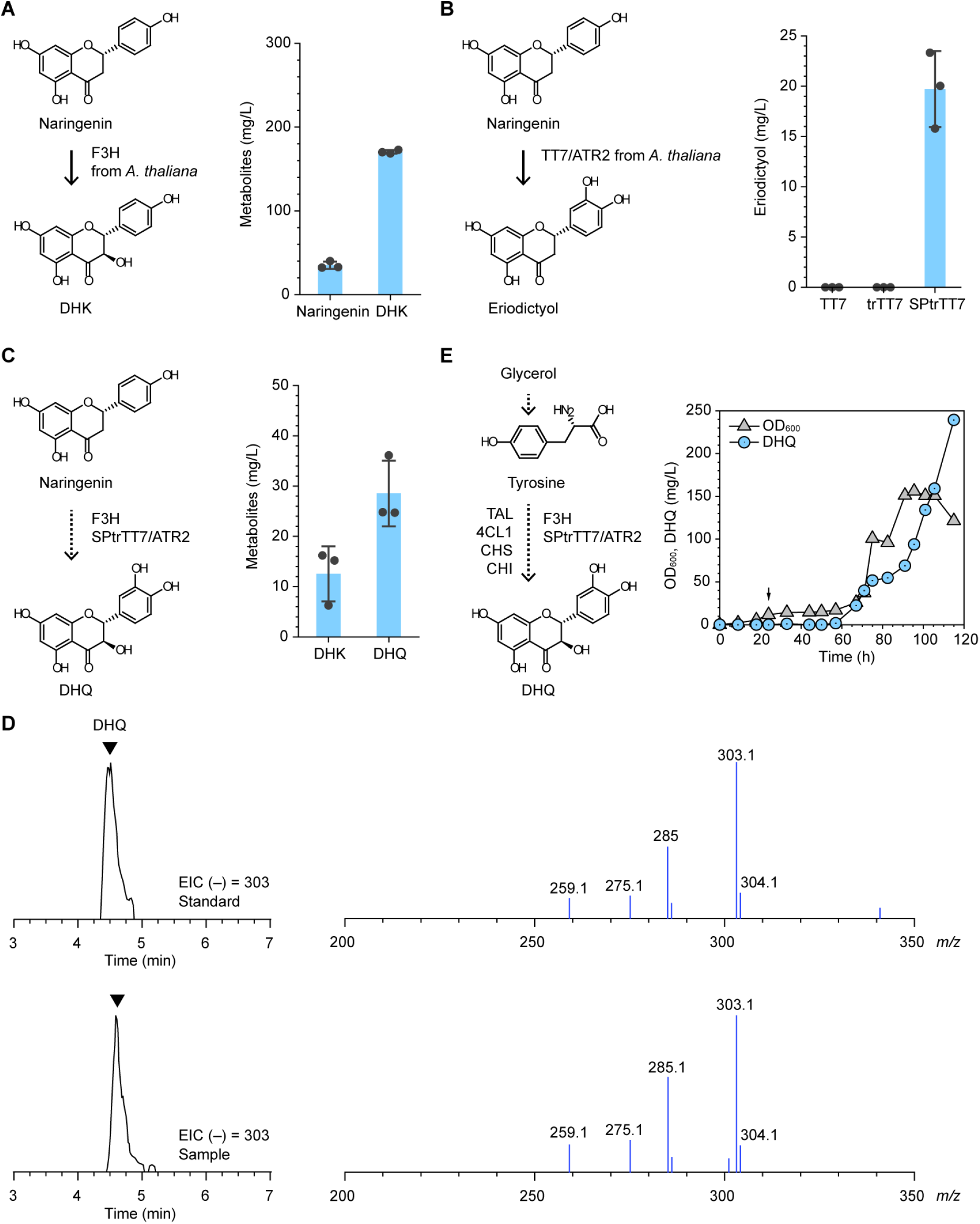
Production of DHQ. **(A)** Conversion of naringenin to DHK by F3H from *A. thaliana*. **(B)** Engineering TT7 for conversion of naringenin to eriodictyol. trTT7, a TT7 variant with its N-terminal signal peptide truncated; SPtrTT7, a TT7 variant with its N-terminal signal peptide exchanged with that of *E. coli* OmpA. **(C)** Conversion of naringenin to DHQ by F3H and SPtrTT7. Residual DHK is also shown. **(D)** Fed-batch fermentation profile of the DHQ strain in R/2 medium. Arrow in the graph indicates IPTG induction time point. (E) Extracted ion chromatograms (EICs) and mass spectrums of DHQ generated by LC-MS, from the authentic standard (upper panels) or from the culture sample of the DHQ strain (lower panels). Data are representative of three independent replicates. **(A-C)** Error bars, mean ± SD (*n* = 3).

Production of DHQ from naringenin can be achieved via either eriodictyol or dihydrokaempferol (DHK) (Fig. 1). Conversion of naringenin to DHK was first tested. Flavanone 3-hydroxylase (F3H) from *A. thaliana* was reported to convert either naringenin into DHK or eriodictyol into DHQ (Owens et al., 2008). Introduction of F3H into *E. coli* BL21(DE3) resulted in 170.4 mg/L of DHK production from 500 mg/L of naringenin (Fig. 1A). Next, a flavonoid 3’-hydroxylase (TT7) from *A. thaliana*, which is a cytochrome P450 (simply P450 henceforth), was employed to convert naringenin into eriodictyol or DHK into DHQ (Schoenbohm et al., 2000). As a P450 monooxygenase requires co-expression of a P450 reductase as its redox partner, a P450 reductase from *A. thaliana* (ATR2) was also employed (Niu et al., 2017). Plasmids pTrc99A-TT7 and pTac15K-ATR2 were constructed and transformed into *E. coli* BL21(DE3), followed by flask culture. However, conversion of naringenin into eriodictyol was not observed (Fig. 1B). By contrast to bacterial P450s, plant P450s are bound to subcellular organelles [i.e., endoplasmic reticulum (ER)], guided by N-terminal signal peptides. In this regard, a TT7 variant truncated with its N-terminal signal peptide (trTT7) was constructed for cytoplasmic expression in *E. coli*. To mimic the mode of action of plant P450s, we also tested guiding the enzyme to the inner membrane by construction of SPtrTT7 (a TT7 variant with its N-terminal signal peptide exchanged with that of *E. coli* OmpA). Activity of the P450 enzyme was only observed when SPtrTT7 was employed, producing 19.7 mg/L of eriodictyol from 100 mg/L of naringenin (Fig. 1B). Such drastic enhancement of enzyme activity can be attributed to two reasons as follows: (1) the membrane-bound SPtrTT7 and ATR2 would have been properly oriented which is favorable for the enzymatic reactions, (2) exchanging the N-terminal tag enhanced enzyme expression as identified by SDS-PAGE analysis (Fig. 2A). Next, to test whether co-expression of F3H and SPtrTT7 would lead to the production of DHQ from naringenin, pTac15K-F3H and pDHQ (harboring *ATR2* and *SPtrTT7*) were introduced into *E. coli* BL21(DE3), successfully producing 28.5 mg/L of DHQ from 100 mg/L of naringenin (Fig. 1C).

**Fig. 2.**
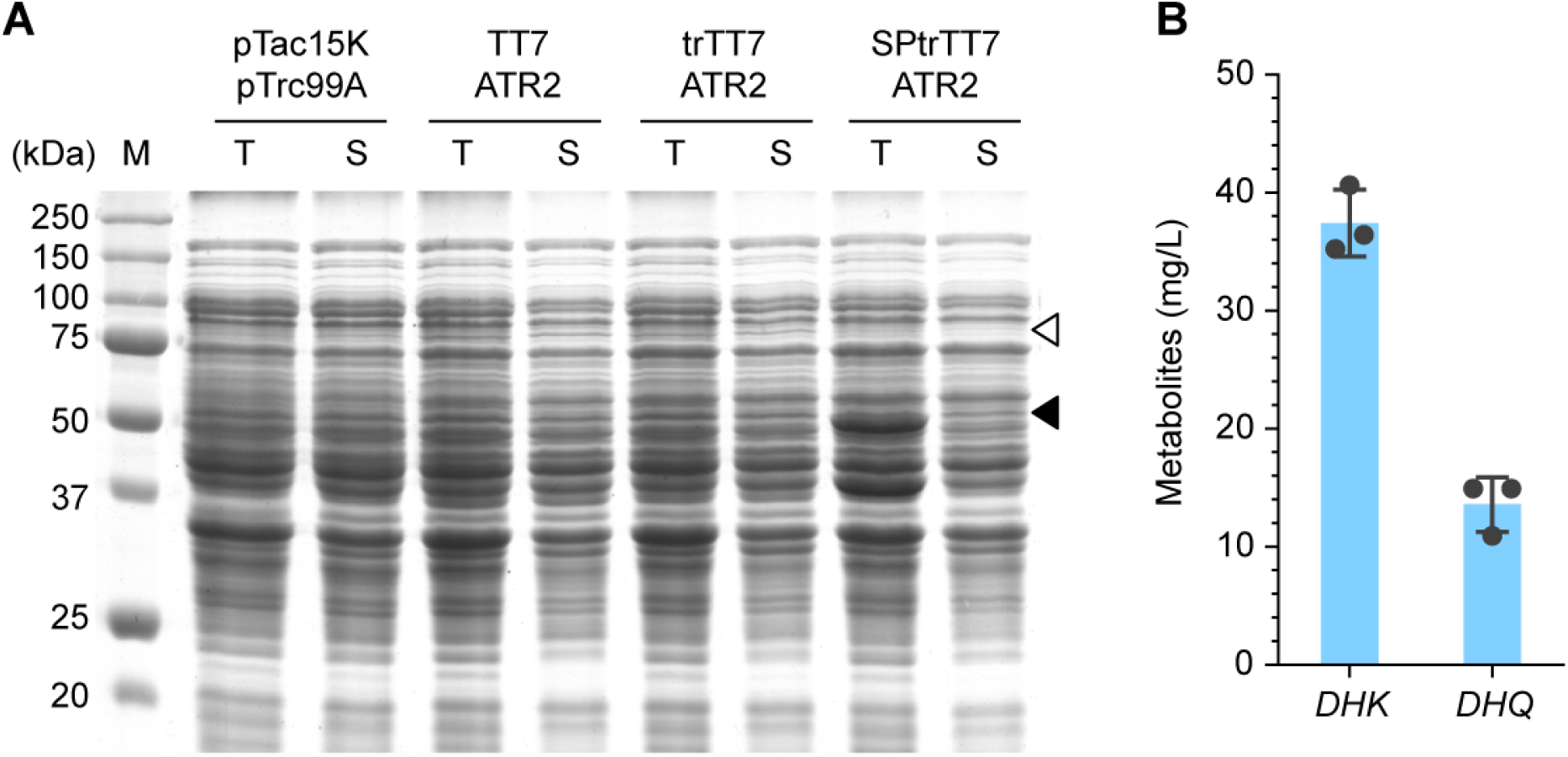
Production of DHQ. (A) SDS-PAGE analysis of *E. coli* BL21(DE3) strains harboring pTac15K-derived plasmids harboring the noted TT7 variants (pTac15K denotes the control without TT7) and pTrc99A-derived plasmids harboring ATR2 (pTrc99A denotes the control without ATR2). T, total fraction; S, soluble fraction; M, protein size marker. Empty triangle denotes ATR2 expression; black triangle denotes TT7 variants expression. (B) Conversion of naringenin to DHQ when *hpaBC* was employed instead of TT7 and ATR2. Residual DHK is also shown. Error bars, mean ± SD (*n* = 3).

To test a non-P450 enzyme—which was expected to be more favorable than P450 in *E. coli—*for DHQ production, *E. coli hpaBC* was tested for its ability to convert naringenin to eriodictyol (Jones et al., 2016b). *E. coli* BL21(DE3) harboring pTac15K-F3H and pCAF (harboring *E. coli hpaBC*) was cultured, and 13.6 mg/L of DHQ was produced from 100 mg/L of naringenin (Fig. 2B). The DHQ titer (13.6 mg/L) was significantly lower than that (28.5 mg/L) obtained by employing SPtrTT7. In addition, higher accumulation DHK (37.4 mg/L) than that obtained by employing SPtrTT7 (12.6 mg/L) revealed that HpaBC has lower conversion efficiency than that of SptrTT7 (Fig. 2B). Since plant P450s are generally difficult to be functionally expressed in bacterial hosts such as *E. coli*, these results suggest that engineering the signal peptide of plant P450s can be an effective solution for bacterial production of diverse natural products requiring plant P450s.

To complete the *de novo* biosynthetic pathway for DHQ production, the *F3H* gene was inserted into the previously constructed pNRG plasmid, resulting in pDHK. Plasmids pDHK and pDHQ were introduced into the *p*-coumaric acid producer (BTY5 harboring pCOU), resulting in the DHQ strain. Shake flask culture of the DHQ strain in MR medium supplemented with 20 g/L of glycerol and 2 g/L of yeast extract resulted in 20.1 mg/L of DHQ production. Glycerol was used as the carbon source because higher production of naringenin, the precursor to DHQ, was achieved from glycerol when compared to that achieved from glucose in our previous study (Yang et al., 2018). Production of DHQ was confirmed by LC-MS analysis (Fig. 1D).

Next, fed-batch fermentation was performed to test the performance of the DHQ strain in bioreactors. The DHQ strain was fed-batch cultured in R/2 medium supplemented with 20 g/L of glycerol and 2 g/L of yeast extract, achieving 239.4 mg/L of titer in 115 h (Fig. 1E). This represents the first demonstration of *de novo* production of DHQ from simple carbon sources in *E. coli*.

## 4. Conclusions

In this study, an *E. coli* strain capable of producing an important phenylpropanoid DHQ from glycerol was developed. The DHQ strain was developed based on the previously constructed *p*-coumaric acid overproducer, achieving 239.4 mg/L of DHQ production after optimizing the biosynthetic pathway and engineering the N-terminal signal peptide of a plant-derived P450. To the best of our knowledge, this is the first report on one-step production of DHQ in *E. coli*. These results also represent the highest titer of DHQ obtained in *E. coli*. Although higher titers might be obtained by stepwise culture, switching from a multistep process (stepwise culture) to a single step process (one-step *de novo* production) is important. This is because employing a single microbial strain is much simpler, making the process more economical, less laborious, and time-saving. This would lead to higher overall yield and productivity. The metabolic engineering and process optimization strategies employed in this study is generally applicable for the production of complex natural products, particularly those requiring multiple precursors. Microbial cell factories developed using such strategies will enable robust and sustainable production of important natural products, significantly contributing to chemical, cosmetic, pharmaceutical, and food industries in the next years to come.

## Acknowledgements

This work was supported by the Technology Development Program to Solve Climate Changes on Systems Metabolic Engineering for Biorefineries [NRF-2012M1A2A2026556 and NRF-2012M1A2A2026557] from the Ministry of Science and ICT through the National Research Foundation of Korea.

## Author contributions

S.Y.L. conceived the project, S.Y.P., D.Y., S.H.H., and S.Y.L. designed the experiments. S.Y.P., D.Y., and S.H.H. conducted the experiments. S.Y.P. and D.Y. analyzed the data. S.Y.P., D.Y., and S.Y.L. wrote the manuscript. All authors read and approved the final manuscript.

